# Structural Characterization of DNA Binding Domain of Essential Mammalian Protein TTF 1

**DOI:** 10.1101/2024.07.05.602234

**Authors:** Gajender Singh, Abhinetra Jagdish Bhopale, Saloni Khatri, Prashant Prakash, Rajnish Kumar, Sukh Mahendra Singh, Samarendra Kumar Singh

## Abstract

Transcription termination factor 1 (TTF1), being multifunctional in nature, is involved in a wide range of critical processes that make it essential for survival of the mammalian cells. TTF1 protein comprises three functional domains: the N-terminal (regulatory/inhibitory) domain, trans-activation domain, and C-terminal domain. The Myb domain is responsible for DNA-binding function of this protein and spans 550 to 732 amino acids (183 residues long). Despite the essential role of TTF1 in multiple cellular processes, there is no physical structure available to date. Purification of the functional full-length protein has been unsuccessful so far. Hence, we moved forward towards characterizing the Myb domain of this essential protein. We first constructed a three-dimensional model of the Myb domain using Robetta server and determined its stability through MD simulation in an explicit solvent. To validate the model, upon codon optimization we cloned this domain into a bacterial expression vector. The protein was then purified to homogeneity and its DNA-binding activity was checked by electro-mobility shift assay. We then proceeded to CD spectroscopy and Raman spectroscopy for secondary structure characterization. The results validated the computational model, concluding that this domain is predominantly helical in nature. The confidence built by this study now pushes us to move ahead in order to solve the atomic structure of this critical domain by crystallography or NMR spectroscopy, which in turn will decipher the exact mechanism by which this essential protein engages DNA to cater to various functions.

## Introduction

The eukaryotic genome possesses multiple copies of rDNA, which are present in tandem and repetitive sequences that encode pre-rRNA and regulatory elements. These regulatory elements allow site-specific termination of RNA Pol I transcription from yeast to humans. RNA polymerase I (Pol I) mediated transcription is terminated in a polar manner by specialized transcription terminator proteins binding to specific sites, which also prevents the collision of replication forks heading in opposite directions. Terminator proteins that mediate transcription termination have been found in a variety of taxa which includes mammalian TTF1, Nsi1 [also known as yeast transcription terminator1 (Ytt1)] of *S. cerevisiae*, Reb1 (RNA polymerase I enhancer binding protein) of *Schizosaccharomyces pombe*, and Rib2 of Xenopus (Jaiswal et al., 2016)

Transcription Termination Factor 1 (TTF1) is a multifunctional nucleolar DNA-binding protein that is involved in various processes, such as Pol I-mediated transcription initiation (Németh et al., 2008) and termination (Grummt et al., 1986), pre-rRNA processing, chromatin remodelling, DNA damage repair (Tiwari. K. et al., 2023), and polar replication fork arrest (Németh et al., 2013). The TTF1 gene is located at the 2; 2 A3 location on mouse chromosome 2 and at 9q34.13, on the long arm of human chromosome 9. This factor binds in an orientation-dependent manner to the terminator element called Sal box, which is an 18-bp sequence motif-**AGGTCGACCAGA/TT/ANTCCG** in mouse and 11-bp long sequence – **GGGTCGACCAG** in humans. TTF1 binding sites are present both upstream and downstream of the rDNA coding region, which brings terminator and promoter loci in close proximity upon TTF1 binding. This allows the direct transfer of Pol I machinery from 3’ end terminator to 5’ end promoter of the adjacent rDNA unit. (Ribomoter model, Goodfellow & Zomerdijk, 2013).

TTF1 protein consists of three functional domains: 1) an N-terminal regulatory domain (NRD), 2) a trans-activation domain (TAD) or central domain, and 3) a C-terminal DNA-binding domain (Figure 1) (Sander & Grummt, 1997). The N-terminal domain (less conserved region between murine and human) is an auto-regulatory domain that is involved in the oligomerization of protein which masks the DNA-binding activity of TTF1. A specific region ranging from 323 to 445 amino acids in the central trans-activation domain is involved in chromatin-specific functions, including transcription termination, chromatin remodelling, and transcription activation. Additionally, another region ranging from 430 to 445 amino acids is crucial for termination, as its deletion affects termination without compromising its DNA-binding activity (Evers et al., 1995). The C-terminal half of the TTF1 protein exhibits a highly conserved amino acid sequence in both mouse and human (Everst & Grummt, 1995). It has two conserved Myb/SANT (**S**WI3-**A**DA2-**N**-CoR-**T**FIIIB) like domains that are involved in DNA binding. This domain shows strong homology with the Reb1 protein of *S. pombe* and the DNA-binding domain of the proto-onco-protein c-Myb. It contains clustered conserved tryptophan residues homologous to Reb1p and c-Myb, which are essential for DNA-protein interactions (Evers et al., 1995). The last 31 amino acids at the C-terminal end of the TTF1 protein are known to show species specificity (Németh et al., 2013).

**Figure 1:**
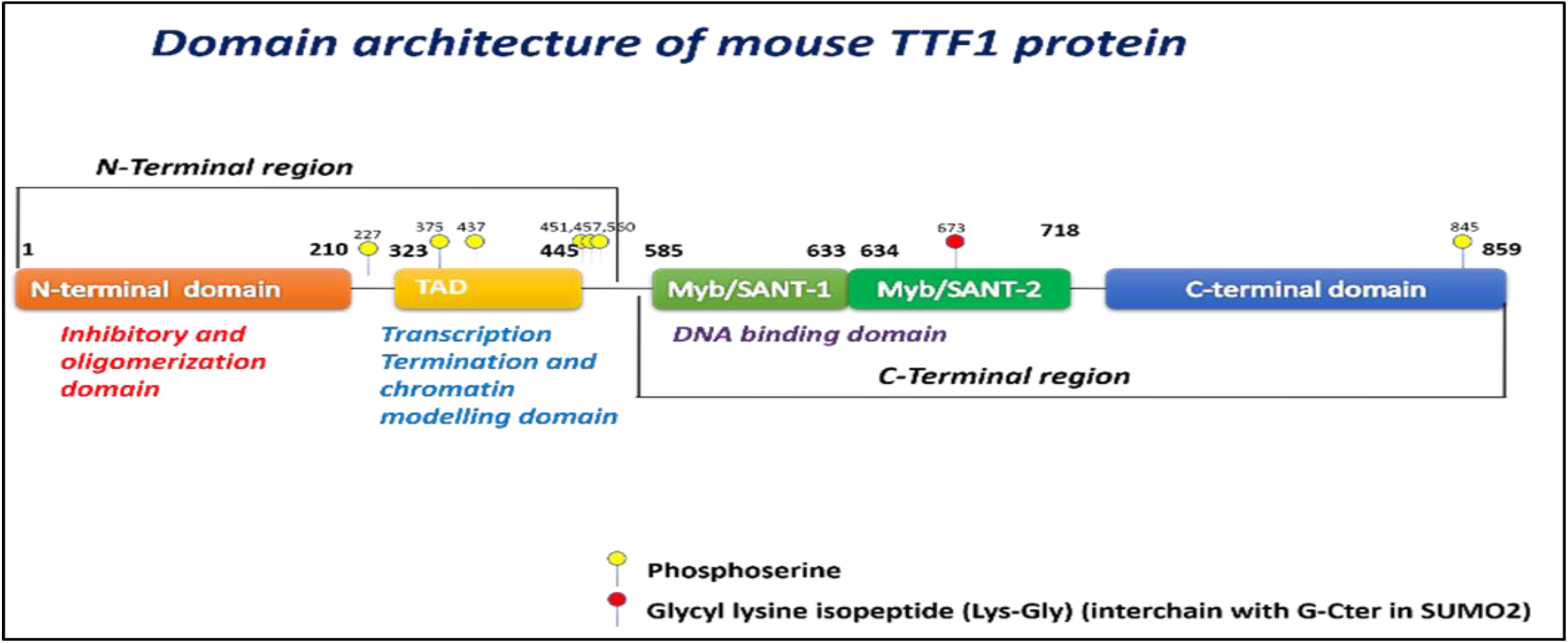
Domain architecture of mouse TTF1 protein showing the N-terminal inhibitory and oligomerization domain (1-210 aa), transcription termination and chromatin remodelling domain (323–445 aa), conserved DNA-binding domain (Myb/SANT –like domain, 550-718 aa), and C-terminal domain (732–859 aa).

Several studies have demonstrated that different domains of TTF1 interact with various important factors that are involved in numerous regulatory processes. Some important TTF1 interacting protein are Cockayne Syndrome group B (CSB), Murine Double Minute 2 (MDM2), Alternative Reading Frame (ARF p19^ARF^ in mouse and p14^ARF^ in human), TTF1 interacting protein 5 (Tip5), Nucleosome remodelling and histone deacetylase complex (NuRD), p300/CBP associated factor (PCAF), Polymerase 1 and transcript release factor (PTRF), and nucleophosmin NPM/ B23. Our lab has recently identified DDB1 as a novel interacting partner of TTF1 and its role in UV-mediated DNA damage sensing (Németh et al., 2013;Tiwari. K. et al., 2023)The C-terminal region, besides being highly conserved, plays an important role in DNA binding and interacts with various proteins to perform the essential functions required for cell growth and proliferation.

It is crucial to explore the structural aspects of these domains to examine the specific amino acid residues engaged in the interaction with other partners. However, neither an *in silico* nor a physically determined structure of the individual domains of TTF1 is available to date. Since the Myb domain of TTF1 is critical for coordinating many cellular processes, in the current study, we predicted the *in silico* model of the Myb domain of mouse TTF1 protein using *ab-initio* and homology modelling and validated the same experimentally as well. We determined the stability of the structure through a 200 ns molecular dynamics (MD) simulation (in triplicate) in an explicit solvent. Further, for physical characterization, we cloned and expressed the Myb domain of the mouse TTF1 protein; we then confirmed its secondary structure using circular dichroism (CD) and Raman spectroscopy. The computational and biophysical analyses of the Myb domain will provide insight into uncovering the atomic structure, which intern will elucidate the mechanism its diverse function.

## MATERIALS AND METHODS

### 1. Sequence retrieval and sequence analysis

The Myb Domain sequence of *Mus musculus* TTF1 was retrieved from UniProtKB (Accession Number: Q62187), in the FASTA format (supplementary, Box 1). The physicochemical features of the Myb domain were observed and compared using Expasy ProtParam (https://web.expasy.org/protparam/). The molecular weight, instability index, isoelectric point (pI), and grand average of hydropathicity (GRAVY) values corresponding to each domain were compared. To evaluate the sequence of the Myb domain, the disorder profile was analyzed using the DisoPred3 web server (http://bioinf.cs.ucl.ac.uk/psipred/).

### 2. Molecular modelling of Myb domain of TTF1, refinement and validation

The DNA binding domain (Myb domain) of the TTF1 protein is widely recognized for its ability to engage with r-DNA’s Sal box terminator elements to stop Pol I-mediated transcription. A sequence homology search was performed for the Myb domain of the mouse TTF1 protein in PDB using BLASTp. For the structural analysis we have generated computational models of Myb domain by the Alpha Fold (https://alphafold.ebi.ac.uk/. Jumper et al., 2021; Varadi et al., 2022), and SWISSMODEL (https://swissmodel.expasy.org/, Homology-modelling server, (Bienert et al., 2017; Guex et al., 2009; Waterhouse et al., 2018). Further, *ab-initio* structure prediction has also been used to obtain the protein structure using I-TASSER (https://zhanggroup.org/I-TASSER/, J. Yang & Zhang, 2015). The Robetta server (https://robetta.bakerlab.org/, Kim et al., 2004) uses both *ab-initio* and comparative modelling techniques to build structural models depending upon the availability of a suitable template structure for homology modelling. The predicted model structures were refined using ModRefiner server (https://zhanggroup.org/ModRefiner/). The structural quality of these models was checked using the SAVESv6.0 web server (https://saves.mbi.ucla.edu/), and the refined model was validated via ERRAT, VERIFY3D, PROCHECK, and Ramachandran plot. The quality of the modelled protein structures was analyzed for Z-score using the ProSA server (https://prosa.services.came.sbg.ac.at/prosa.php), and the percentages of the secondary structures were determined using VADAR version 1.8 (http://vadar.wishartlab.com, Willard et al., 2003).

### 3. Molecular Dynamics Simulation

Molecular dynamics study was performed using GROMACS 2020 and CHARMM36 force field (Smith et al., 2015). The Myb domain was solvated in a cubic box using a TIP3 water model with a distance of 1 nm between each side and ionized with Na^+^ and Cl^-^ ions at a concentration of 0.15M to neutralize the system. The constructed system contained 66 Na^+^ ions, 85 Cl^-^ ions and 23190 water molecules. Furthermore, the system was minimized using the steepest descent algorithm until the maximum force decreased below 1000kJ/mol. During the equilibrium phase, we implemented position restraints under constant number, volume, and temperature (NVT) and isothermal-isobaric (NPT) ensembles for 1 ns each to prevent potential distortions that could lead to instability. V-rescale (Bussi et al., 2007) temperature coupling and Parrinello-Rahman pressure coupling (Parrinello & Rahman, 1981) were employed to maintain the system at 300 K temperature and 1 bar pressure along with coupling constant of 0.1 picosecond for temperature and 2 picosecond for pressure. Long-range electrostatic interactions and van der Waals interactions were calculated using the Particle Mesh Ewald method (Essmann et al., 1995), and the cut-off distance for short-range van der Waals interactions was set to 1 nm. The LINCS algorithm (Hess et al., 1997) was used to constrain all bonds. Finally, a 200 ns production simulation was conducted in triplicate using periodic boundary conditions. Furthermore, the final production simulation trajectory was analyzed by calculating the root mean square deviation (RMSD), root mean square fluctuation (RMSF), radius of gyration (RoG), and solvent-accessible surface area.

### 4. Plasmid Constructs

The codon-optimized Myb domain sequence of the TTF1 was amplified using the mentioned primers and cloned into the pET28a expression vector (Novagen, Millipore, USA) using EcoRI and Sal-I restriction sites. The clone was verified by double-restriction digestion and sequencing. For expression, the verified clone was transformed into *E.coli* BL21 (DE3) strain.

Primer Forward: 5’ ACCCTGATTACCAATCTGAAACGC 3’ Primer Reverse: 5’ TTAAACACCACGATAAACAAAACC 3’

### 5. Expression and purification of the TTF1 Myb domain

Myb/pET28a clone vector was transformed into BL21 (λDE3) *E. coli* strain for expression of the recombinant protein and grown in LB culture media at 37 ^°^C at 210 rpm in shaker incubator with 30 µg/ml kanamycin antibiotic until the OD_600_ reaches 0.6 to 0.8. Exponentially grown culture was induced with 0.9 mM isopropyl β-D-1-thiogalactopyranoside (IPTG; HiMedia, India) at 30 ^°^C for approximately 6 to 8 hours. The induced culture was harvested by centrifugation at 8000 rpm for 10 min at 4 ^°^C. The pellet was re-suspended in lysis buffer [25 mM Tris (pH-7.5), 500 mM KCl, 10% (w/v) glycerol, 9 mM β-mercaptoethanol, 10 mM Imidazole, 1mM PMSF, 2 mg/ml lysozyme] with 1X protease inhibitor cocktail (Roche, USA) for 1 hour at 4 ^°^C and subjected to sonication (30 sec pulse with 50 sec intervals for 10 mins at 40% power on ice) to ensure complete lysis. The cell lysate was clarified by centrifugation (14000 rpm, 1 hour at 4 ^°^C) to remove debris. The clear lysate was mixed with pre-equilibrated Ni-NTA beads (Thermo Fisher Scientific, USA) and incubated on a rotatory shaker for 2 h at 4 ^°^C for adequate binding. The mixture was loaded onto a column and allowed to settle under gravity. The packed column was then washed with 10 column volumes of wash buffer [25 mM Tris (pH-7.5), 500 mM KCl, 10% (w/v) glycerol, 9 mM β-mercaptoethanol, 40 mM Imidazole, 1mM PMSF], and the protein was eluted with elution buffer [25 mM Tris (pH-7.5), 500 mM KCl, 10% (w/v) glycerol, 9 mM β-mercaptoethanol, 400 mM Imidazole, 1 mM PMSF]. The eluted protein fractions were pooled and concentrated using a 10 kDa cut-off (Amicon filter, Millipore, USA). After affinity purification, the protein was subjected to size-exclusion gel filtration chromatography using a HiLoad 16/600 Superdex 200 pg preparative SEC column (Cytiva, USA). The protein was resolved on 10% SDS-PAGE at each step from induction to purification in order to observe the induction and level of protein purity. Protein concentration was estimated using the Bradford method, and samples were stored at -80 °C in storage buffer (25 mM Tris (pH-8.0), 300 mM KCl, and 30% glycerol).

### 6. DNA Binding Assay (EMSA)

Reactions were assembled in 20 µl volume containing variable concentrations of protein (TTF1 Myb domain) and 100 ng DNA (Sal Box sequence) were mixed and incubated at room temperature for 40 min in binding buffer (12 mM Tris-HCI, pH 8.0, 100 mM KCl, 5 mM MgCl_2_, 0.1 mM EDTA, and 0.5 mM DTT, 5% glycerol). The protein-DNA complexes were then resolved on a 10% non-denaturing polyacrylamide gel at 4°C and 150V for 2 hours in TBE buffer (Singh et al., 2010). The shifts in DNA bands were visualized using an EMSA kit (Invitrogen, USA) according to the manufacturer’s instruction.

### 7. CD Spectroscopy

The purified Myb domain was diluted to 20 ng/μl in 10 mM Tris-Cl (pH-7.5) and subjected to CD spectroscopy using a Jasco J-1500 spectropolarimeter, having Peltier temperature-controlled cell holders (Jasco, Easton, MD, United States). Data collection and analyses were performed in triplicates (in accordance with Colarusso et al., 2018). Following the standard procedures, the secondary structure was evaluated using the Jasco (J. T. Yang et al., 1986) and OriginPro software (version 2022, Origin Lab Corp., Northampton, MA, United States) according to protocol described by Greenfield el.al. (Greenfield, 2007).

### 8. Raman Spectroscopy

Raman spectroscopy (alpha300, WITec, Germany) was used for recording the Raman spectrum homogeneity of the purified Myb domain. The instrument was equipped with a liquid-nitrogen-cooled charge-coupled device (CCD) detector and a spectrograph with a 600 g/mm grating with a resolution of 1 cm^−1^. For spectrum collection, backscattering geometry was combined with a notch filter to reject elastic contributions. Excitation was achieved using a laser emitting at 532 nm, with a power of 44 mW (Tiwari et al., 2022). Raman spectra of the Myb domain of TTF1 was recoded using a 50 objective with a numerical aperture (NA) ¼ 0.6 (laser spot diameter reaching the sample was about 1 mm). An optical glass slide with a cavity was used for the measurements, which were carried out on sample drops (5 µg/ml) at room temperature. For each sample, five Raman spectra were obtained at an acquisition time of 10 s from various spots on the drops. High signal-to-noise ratio spectra were picked among some of the recorded spectra upon verifying the spectral profile. OriginPro software was used to map and evaluate the same (Signorelli et al., 2017). Protocol as described by Maiti et al., 2004 was used for curve fitting of the amide I band and its assignment to the structural components using the Origin software (Tiwari et al., 2023).

## Results

### 1. Amino acid sequence-based analysis

The amino acid sequence-based physicochemical properties of the Myb domain were computed using the ProtParam server, as listed in Table 1. The isoelectronic point (pI) value of this protein was above 7, suggesting the basic nature of the protein due to a preponderance of basic amino acid residues in its side chain. An instability index of < 40 indicates protein stability. The instability index of 30.86 for the Myb domain, suggest that the domain is stable. This is corroborated by the disorder prediction profile, which shows that the Myb domain is stable when the disorder is less than the threshold value of 0.5 (Figure 2). The hydrophobicity of the protein was determined using the GRAVY method, which assesses the ratio of the total number of residues in the protein sequence to the sum of the hydropathy values of the amino acids. The Myb domain exhibits hydrophilic properties, as evidenced by a negative GRAVY value of -0.672.

**Figure 2:**
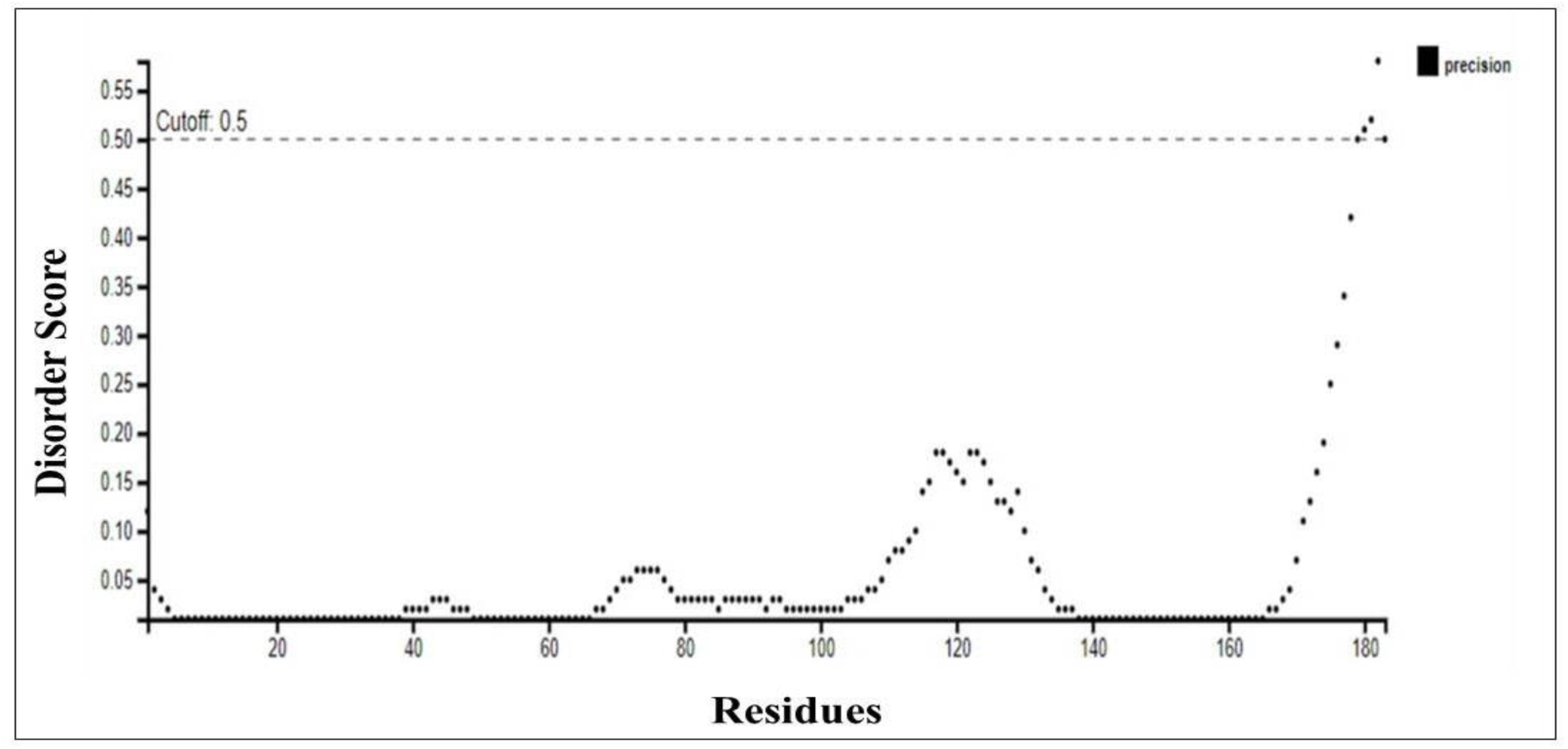
Graphical representation of disordered Myb domain regions in mouse TTF1 protein. Residues with disorder score ≥ 0.5 were considered to be disordered.

**Table 1:** Physicochemical properties of Myb domain of TTF1 protein using ExPASy’s ProtParam tool.

| Physicochemical Properties | Values |
| --- | --- |
| Number of residues | 183 |
| Molecular weight | 21334.80 Da |
| Theoretical pI | 10.11 |
| Molecular formula | C<sub>958</sub>H<sub>1542</sub>N<sub>280</sub>O<sub>262</sub>S<sub>5</sub> |
| Instability index | 30.86 |
| Aliphatic index | 82.57 |
| Number of negatively charged residues (D+E) | 19 |
| Number of positively charged residues (R+K) | 38 |
| Extinction coefficient | 43430 |
| Grand average of hydrophobicity (GRAVY) | -0.672 |
| Estimated half-life (*Escherichia coli*, *in vivo*) | >10hrs |

### 2. Molecular modelling of Myb domain and Validation

To understand the structure of the Myb domain of TTF1, we have predicted computational model of the same. The results of the template search using BLASTp against the Protein Data Bank (PDB) reveal that the highest target-template coverage (Myb domain sequence identity with the template) is 27%, which falls within the ’twilight zone’ for homology modelling (Rost, 1999). For the above reasons, the structure of the Myb domain was modelled using SWISS-MODEL (with the Reb1 protein as a template, PDB ID: 5EYB; Jaiswal et al., 2016) and AlphaFold server (Figure 3A, B). Sequence alignments with its homologue, Reb1 protein, are mentioned in the supplementary file (Figure S1). In addition to homology modelling, a *de-novo* model of the Myb domain was predicted using the Robetta and I-TASSER servers (Figure 3C, D). The models revealed that the Myb domain of TTF1 comprises two Myb-like motifs (illustrated in blue and magenta, Figure 3), which are predominantly composed of alpha-helices. Notably, these motifs are entirely devoid of β-sheets, and the remaining structures are composed of random coils. The structural statistics and different parameters obtained to check the structural reliability using PROCHECK and SAVEv6.0 are provided in Table 2. To assess the quality of the generated structural model, Ramachandran plot analysis was conducted (Figure 3). Marginal variance was observed in the percentage of residues situated within favourably allowed regions for the Myb domain across the AlphaFold, SWISS-MODEL, and Robetta models (Table 2). The residues in the most favoured region in case of all models of the Myb domain were approximately 90%, except for the I-TASSER-predicted Myb model, where 72.1% of residues were present in the most favoured region and 4.2% of residues were present in the disallowed regions. The reason behind above observation could be due to the presence of few residues in the disallowed region for the I-TASSER model, as observed in the Ramachandran plot summary (generated using ProCheck server for structural validation Table 2). The most notable feature among the predicted structural models was the difference in the compactness of the secondary structure. AlphaFold, SWISS-MODEL, and Robetta predicted compact and ordered structures with a very high percentage of α-helical conformations. While the model predicted by I-TASSER was less compact and ordered compared to the models generated by the above servers. All models have very similar and reliable statistics as per the overall SAVESv6.0 results. In summary, the structural integrity and statistics of the models derived from both homology and *ab-initio* methods showed considerable consistency. This confirms the reliability of Robetta and other models (excluding I-TASSER) for further computational analysis. Considering all the above factors, we have selected the Robetta generated *de novo* model of the Myb domain to pursue for further computational analysis. Based on the Z-score (-5.07) of the structure measured using ProSA, it was found that the Myb domain has a structure comparable to other PDB structures of similar size (as determined by NMR, Figure 4A). The quality of the predicted model was highly satisfactory, as evidenced by the fact that the predicted Myb domain structure was superimposed on its yeast homologue Reb1 structure, with an RMSD value of 1.467 nm (Figure 4B). Furthermore, we utilized the VADAR 1.8 server to compute the proportion of secondary structures in the modelled Myb domain. The results indicated that the model consisted of approximately 66.2% α-helices, 0% β-sheets, and 33.2% random coils. These findings support the fact that the majority of DNA-binding proteins are helical in nature in order to stabilize the DNA.

**Figure 3:**
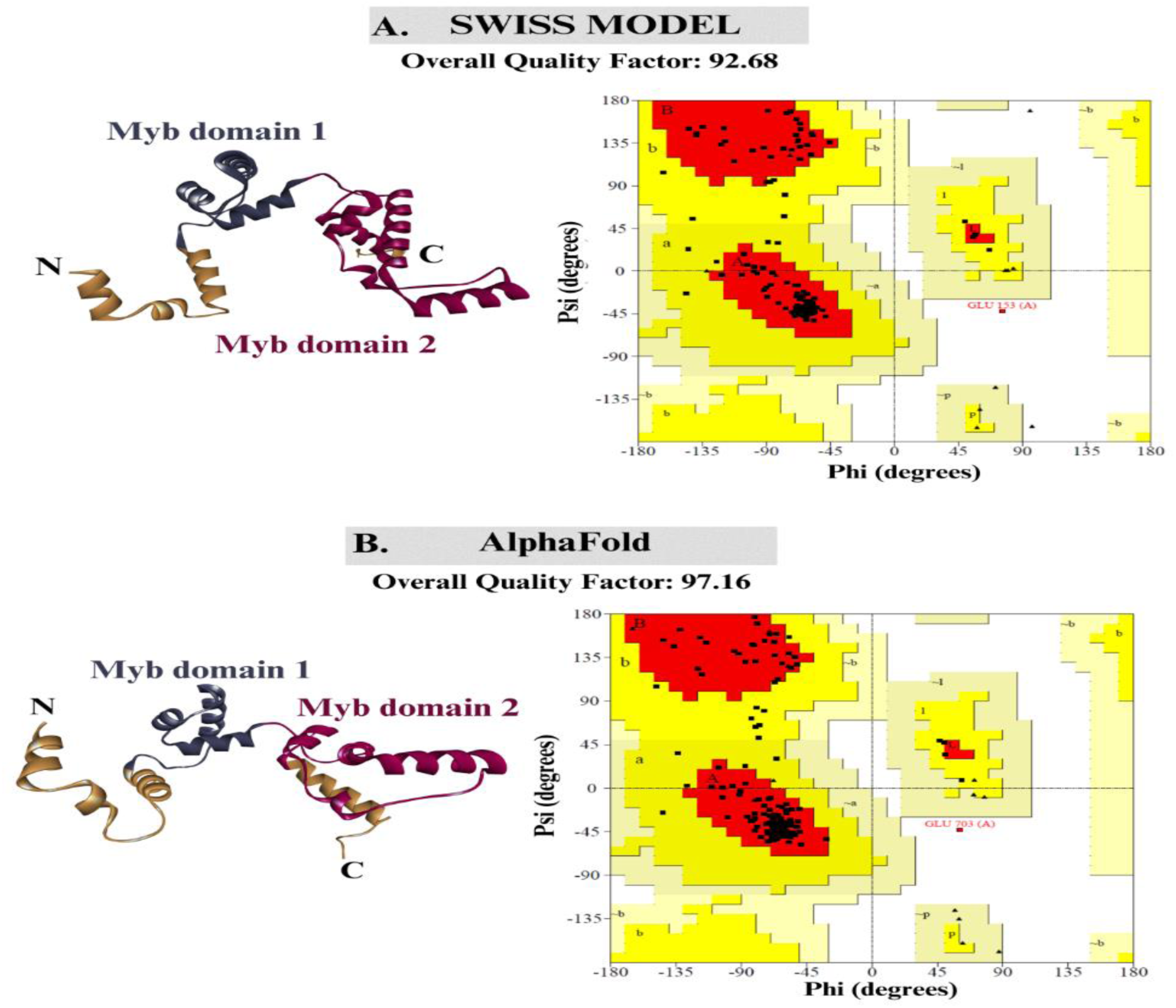
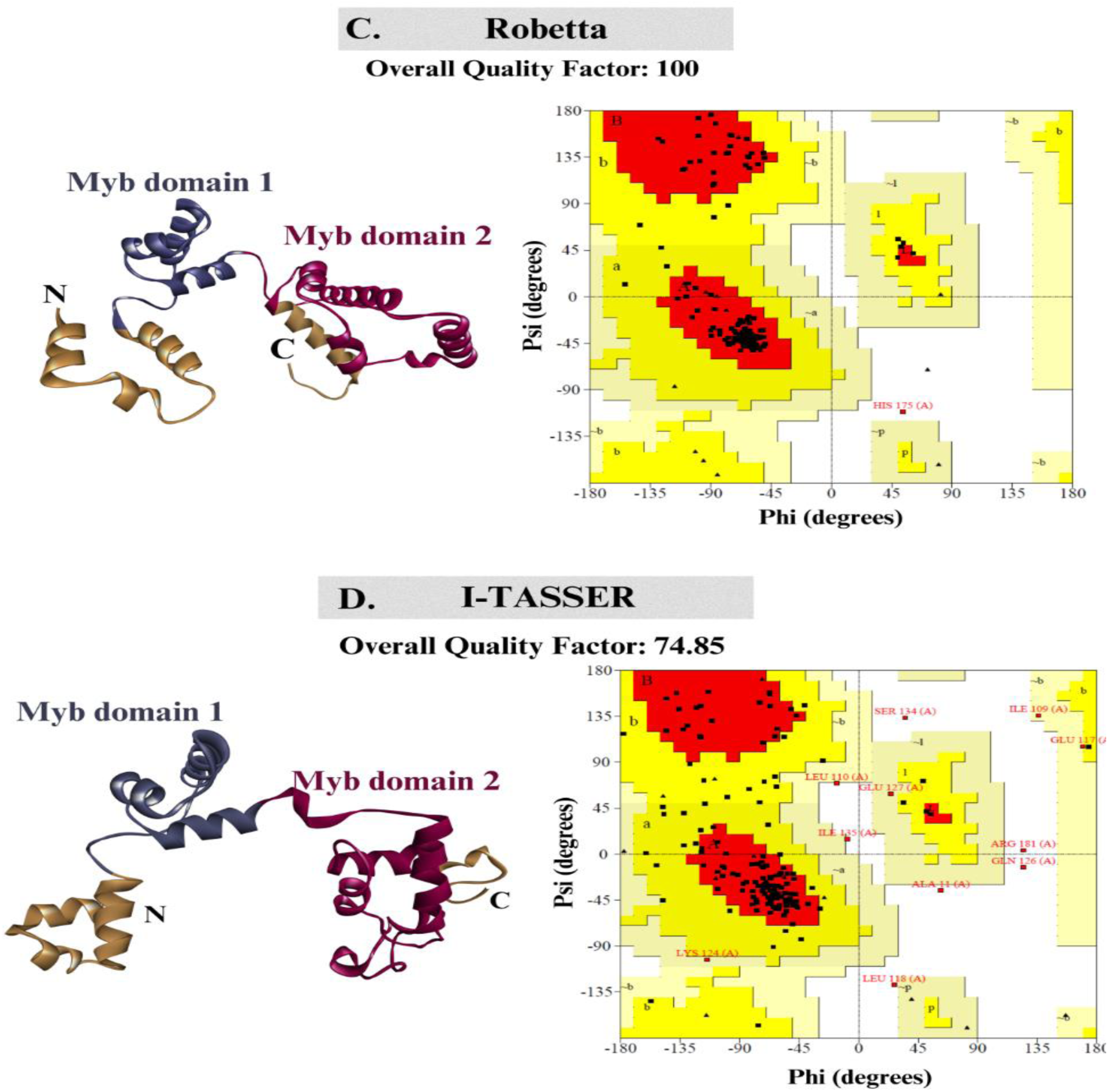
Predicted models of the Myb domain of TTF1 protein by various servers (homology Modelling and *ab-initio* based server) along with Ramachandran plots. The overall quality of the predicted models were analyzed by the SAVEv6.0 (ERRAT) server. ERRAT uses typical nonbonded atomic interactions to distinguish between portions of the structure that were identified properly and erroneously, thereby estimating the overall quality of the proteins.

**Figure 4:**
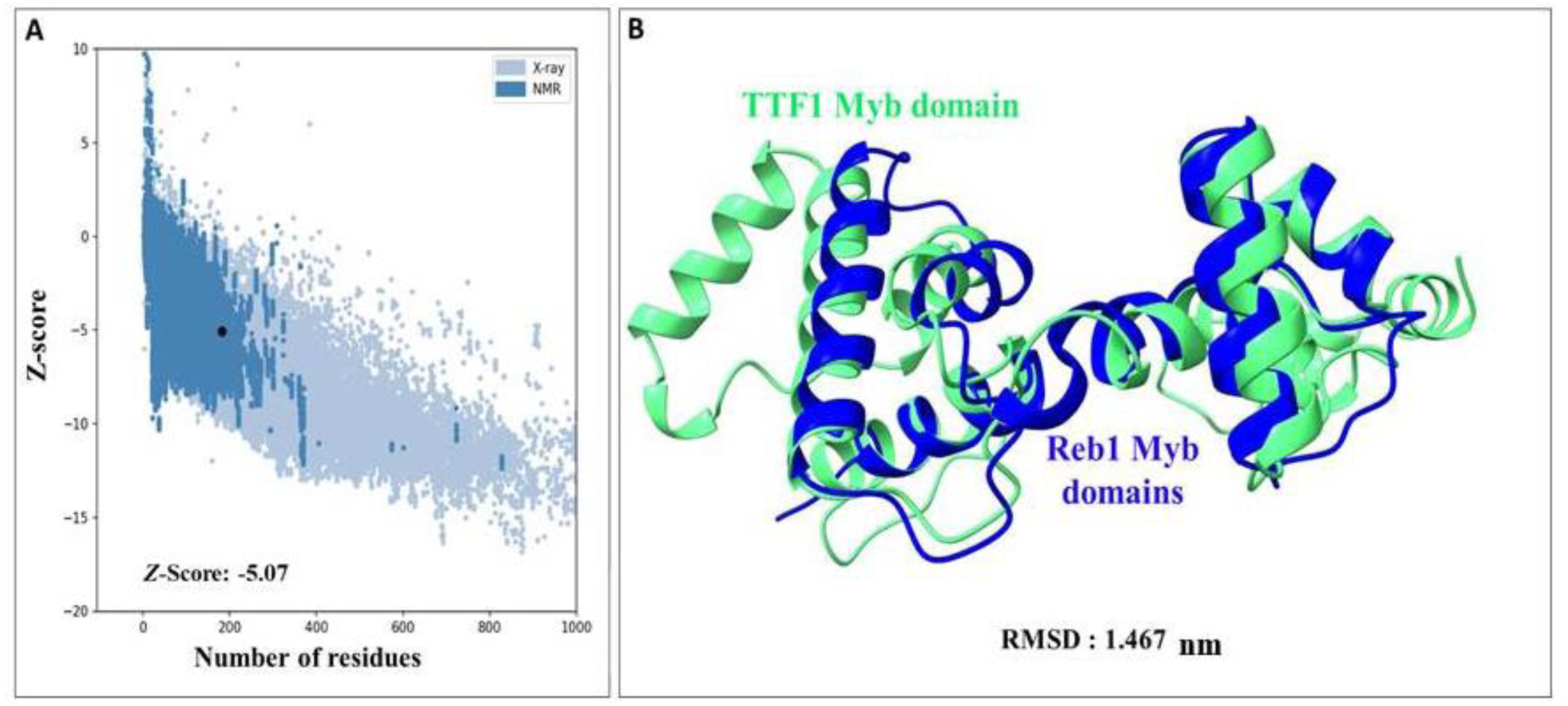
Structure validation of energy-minimized Robetta model of Myb domain by the ProSA server. A) Z – score plot of the Myb domain (represented by black dots ) determined by ProSA. A Z –Score of -5.07, indicates the overall quality of the predicted model. B) Superimposition of the TTF1 Myb domain (green) with the crystal structure of its yeast homologue Reb1 Myb domain (blue).

**Table 2.** The Procheck server assessed the Ramachandran plot data for the anticipated models of the Myb domain.

| PROCHECK Ramachandran plot statistics of Myb domain |  |  |  |  |
| --- | --- | --- | --- | --- |
| Server name | SWISS MODEL | Alphafold | Robetta | I-TASSER |
| Residues in most favored region | 90.9% | 89.1 % | 93.3 % | 72.1% |
| Residues in additional allowed region | 8.5% | 10.3 % | 6.1 % | 21.2% |
| Residues in generously allowed region | 0.0 % | 0.0 % | 0.0 % | 2.4% |
| Residues in disallowed region | 0.6 % | 0.6 % | 0.6 % | 4.2% |

### 3. Trajectory Analyses

To analyze the structural stability and conformational changes in the Myb Domain model obtained by the Robetta server, MD simulation analysis was performed for 200 ns in triplicate. Various statistical parameters, such as root mean square deviation (RMSD), root mean square fluctuation (RMSF), radius of gyration (RoG) (Van Der Spoel et al., 2005), and solvent-accessible surface area, were analyzed from the molecular dynamics (MD) trajectory using the GROMACS analysis tool.

The RMSD was analyzed to determine the deviation from the initial structure, which was calculated by aligning all frames in a trajectory with the first frame throughout the simulation. The RMSD plot indicates that the three runs remained stable throughout the simulation (Figure 5A). The RMSD values were observed within the range of 0.5-2 nm (supplementary figure S2) (for all the three runs). To assess the dynamic behaviour and flexibility of amino acid residues, we analyzed the RMSF of amino acid residues throughout the simulation run. The RMSF plot for the Myb domain of TTF1 across all three runs showed minimal fluctuations, with values ranging from 0.5 to 2 nm. However, the third run exhibited high flexibility in the N-terminal region of the TTF1 Myb domain, with values reaching up to 2.5 nm. This was expected, as the N-terminal region is naturally exposed to the solvent surface (Figure 5B). Furthermore, we performed solvent-accessible surface area (SASA) analysis to determine the surface area of the protein exposed to the fluidic exterior, which helps predict the stability of the hydrophobic core of the protein. Figure 5C shows that the SASA value of the TTF1 Myb domain for all three runs remained stable within the range of 125-155 nm². We also determined the radius of gyration (RoG) of the complex to understand the compactness and overall dimensions of the protein structure. The RoG values for the first two runs were observed within the range of 2–2.5 nm, indicating that the protein maintained a compact, folded conformation throughout the simulation. However, in the third run, the RoG value ranged from 2-3 nm, displaying a minor deviation from the first two simulation trajectories (Figure 5D).

**Figure 5:**
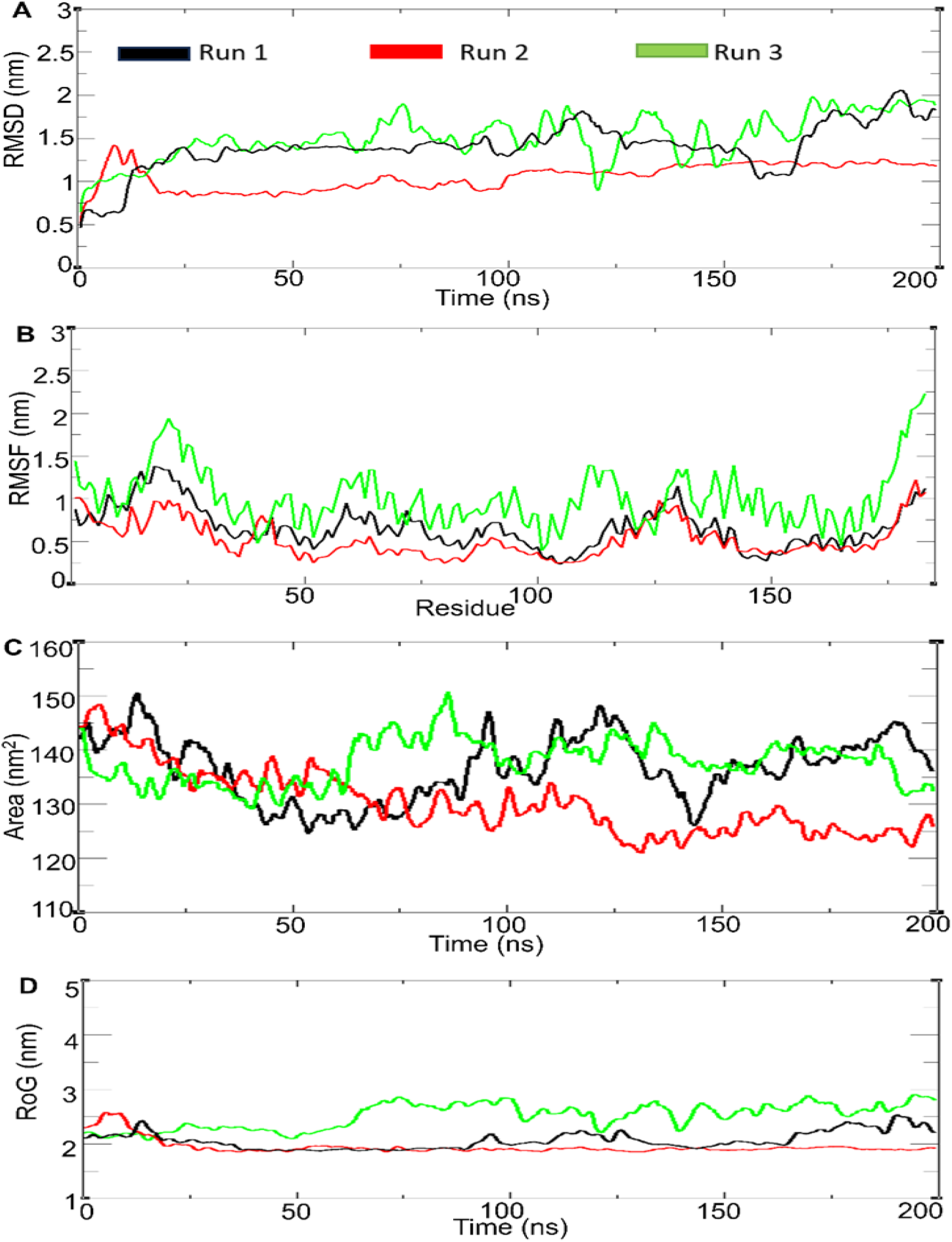
Analysis of MD trajectories of triplicate runs of mammalian protein TTF1. A) Root mean square deviation, B) Root mean square fluctuation, C) Total solvent accessible surface area (y-axis), D) Radius of gyration of simulated protein.

#### 3.1 Free Energy Landscape (FEL) Analysis

FEL analyzes the Gibbs free energy of the system, whereas, in terms of protein conformation, it determines its most stable energy form. It integrates the radius of gyration and the RMSD variables to reflect the specific properties of the system. In the 2D energy landscape, the centralized blue area represents the complex within a cluster with the minimum energy and maximum stability (Figure 6). In 3D projection, the narrow-shaped funnel illustrates the changing conformation of the protein over time, demonstrating dynamic shifts within the system until it settles into a stable structure with low energy (Mallamace et al., 2016; Onuchic et al., 1997). The results showed the changes in the Gibbs free energy (ΔG) values ranging from 0-11.70 kJ/mol for first two run and 0-7.72 kJ/mol for third run. In Figure 6, the formation of a single narrow folding funnel in the 3D plot illustrates a stable folding process for the first two runs. This, along with the 2D contour plot, indicated that it had one local energy minimum, reflecting a stable folding process. However, in the third simulation run, the trajectory diverged from that of the first two, displaying multiple folding patterns and energy minima, indicating an unstable folding process. The 3D conformational state of the TTF1 Myb domain, as determined from the lowest energy minima across the three independent simulation runs are represented in the Figure 6. Overall, the results indicated that the TTF1 Myb domain remained stable throughout the simulation time with high accuracy.

**Figure 6:**
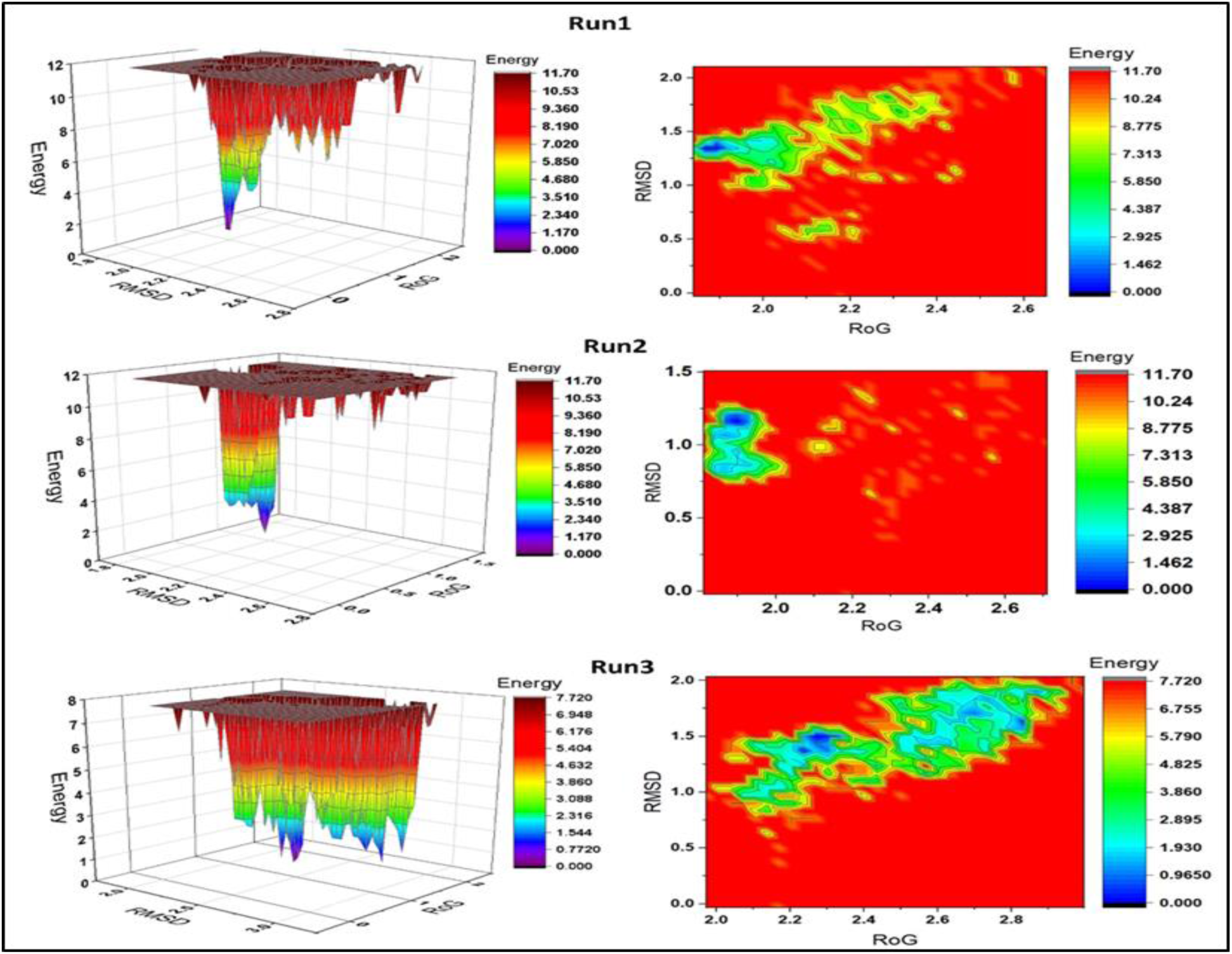
3D and 2D free-energy landscape diagrams as functions of RMSD and RoG as the two coordinates of the TTF1 Myb Domain. Energy is shown in kJ/mol and is indicated by a colored heat map, where deep red indicates the highest energy and deep blue indicates the lowest energy. The 3D projections and 2D contour plots were plotted using the OriginPro2019 software. RMSD: Root mean square deviation, RoG: Radius of gyration.

#### 3.2 Close contact map analysis

Residue-residue close contact map analysis identified amino acid residues that interacted with each other during simulation. Residue-residue close contact maps for the lowest energy conformation of the protein obtained from FEL analysis were created using gmx mdmat, with a cutoff distance of 1.5 nm (Maiorov & Crippen, 1995). The tool calculates the inter-residue distance matrix and provides the average close contact between residues. Amino acid residues 140-170 remain in close contact with residues 80-100 for all three runs (Figure 7). Moreover, for the first two runs, residues 80-90 maintained close interactions with residues 10-60, but these contacts were absent in the third run. Likewise, the close contact between amino acid residues 150-165 and residues 15-25 observed in the first two runs was absent in the third run. This difference might account for the deviation in the results of the third run from those of the first two runs.

**Figure 7:**
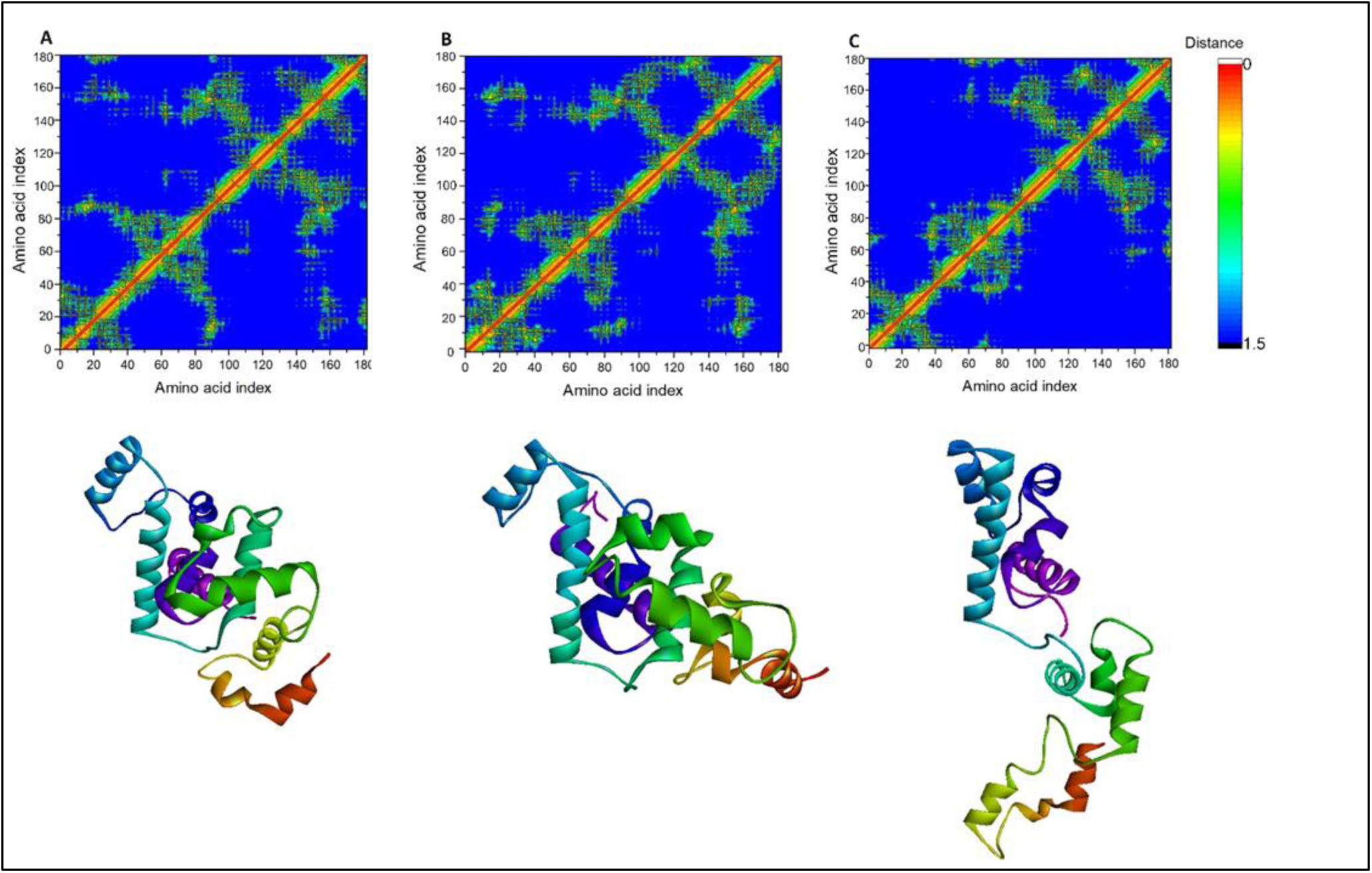
Close contact map and structural representation of the TTF1 Myb domain in its lowest energy state, as identified from Free Energy Landscape (FEL) analysis. The results are shown for the first simulation run (A), second simulation run (B), and third simulation run (C).

Furthermore, to determine the structural stability of the TTF1 Myb domain, we extracted frames from the first simulation run every 20 ns and aligned each extracted frame using the PyMOL tool. Supplementary 3we observed that all frames were aligned with minor deviations and fluctuations, indicating the stability of the TTF1 Myb domain over the simulation time (Figure S2).

### 4. Purification of TTF1 protein Myb domain

The codon-optimized mouse Myb domain of TTF1 was amplified and cloned into the bacterial expression vector pET28a, as described in the Methods section (Figure 8A). The induced protein was purified using Ni-NTA-based affinity chromatography (Figure S3) followed by size exclusion chromatography (as mentioned in the Materials and Methods section). Protein purity was confirmed by SDS-PAGE analysis and homogeneity by Dynamic light scattering (DLS). The purified protein was concentrated to 1 mg/ml (Figure 8B). From the elution profile (Figure 8C) and DLS data (data not shown here), it was confirmed that it was highly homogeneous and mono-dispersed, respectively.

**Figure 8:**
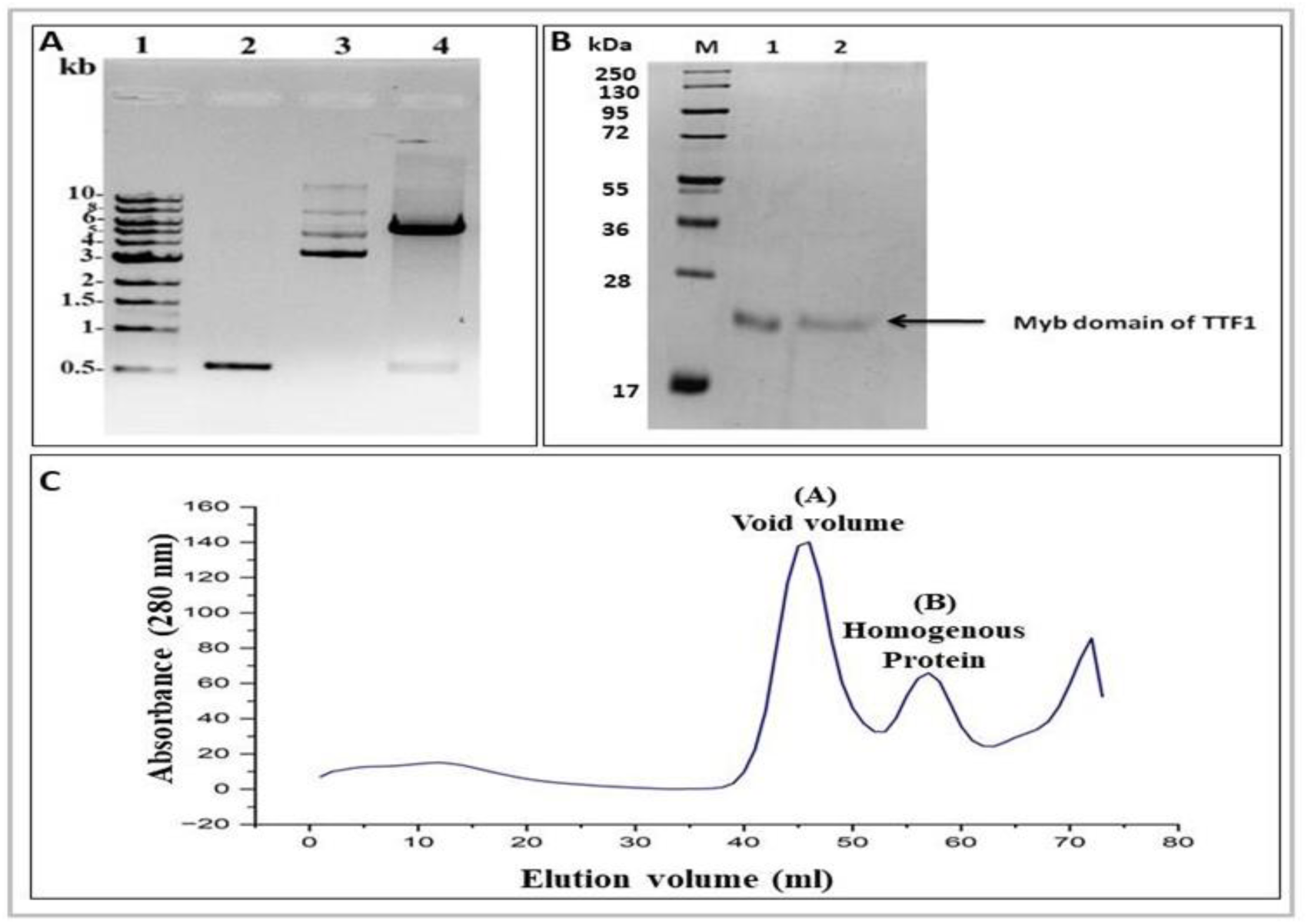
Cloning and purification profile of the Myb domain of mouse TTF1 protein. (A) Agarose gel profile of cloning procedure, Lane 1; DNA Ladder, Lane 2; PCR amplified insert, Lane 3; Cloned vector, Lane 4; Double restriction digested clone vector by *EcoRI* and *Sal I*. (B) Purification profile of the Myb domain protein: Lanes 2-3 show eluted Myb domain. M represents the molecular weight marker. (C) Size exclusion chromatography profile of Myb domain protein on a HiLoad 16/600 Superdex 200 pg preparative SEC column. Peak A represents the void volume, and peak B represents the Myb domain elution profile.

### 5. Electrophoretic mobility shift assay (EMSA)

Myb domains of various transcription factors are involved in DNA-binding activity. Since TTF1 is a transcription factor, we wanted to determine whether the Myb domain alone can bind to DNA. For this, we carried out an electrophoretic mobility shift assay (EMSA), which is a well-established and widely used method to confirm DNA-binding activity. The DNA-protein complex was allowed to form using a fixed quantity (100 ng) of the Sal box DNA sequence and progressively increasing amount of purified Myb domain protein. Upon resolving the protein-DNA complex on a 6% native polyacrylamide gel, it was observed that increasing the amount of protein stoichiometrically shifted the Sal box DNA. The results confirmed that the Myb domain alone was sufficient to bind to the Sal box DNA sequence (Figure 9A).

**Figure 9:**
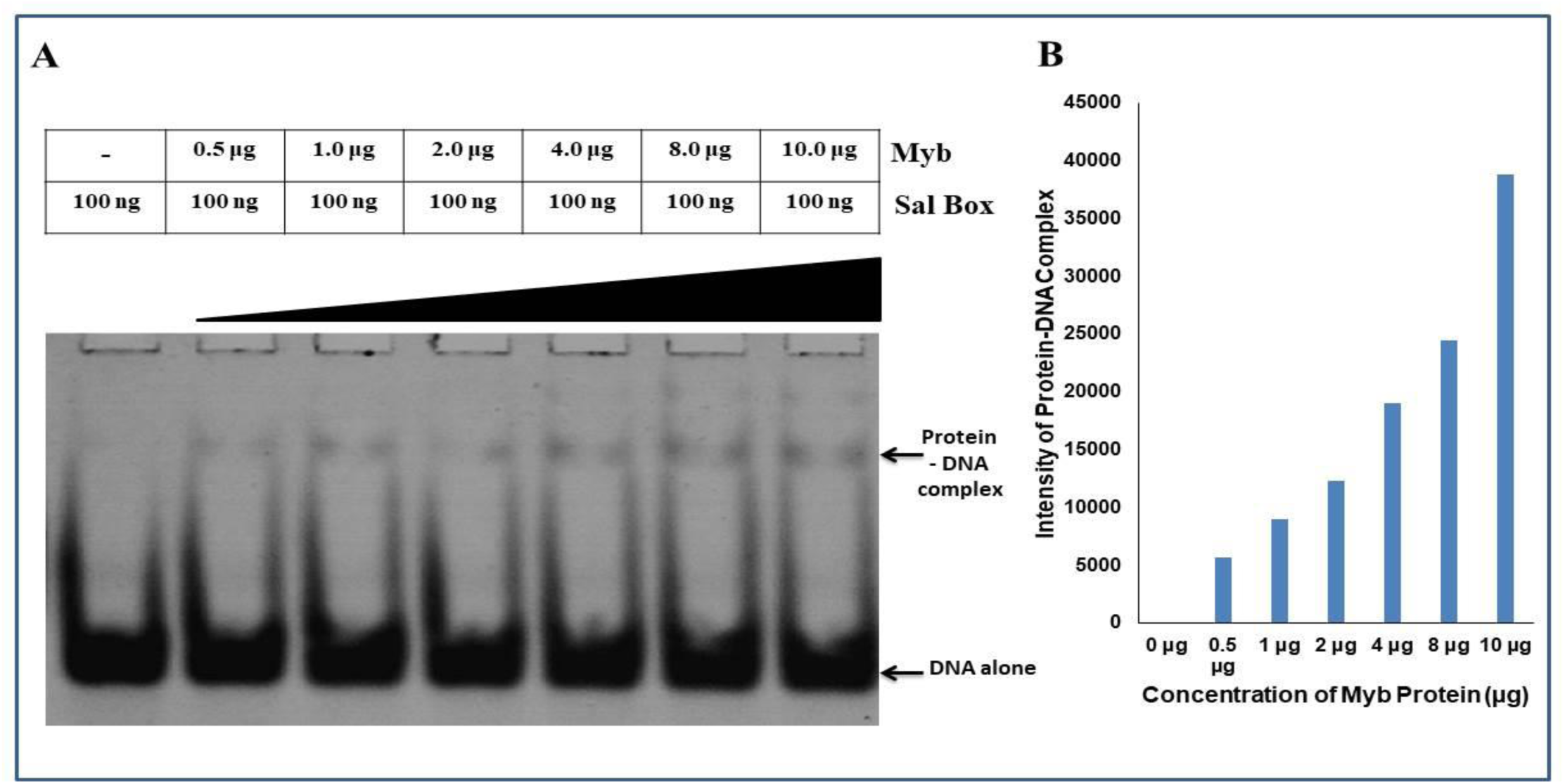
(A) Electrophoretic mobility shift assay (EMSA) to determine the DNA-binding activity of the Myb domain. A shift in the DNA band was detected using SYBR Green (EMSA kit, Invitrogen). Portions of DNA alone and protein-DNA complexes are shown at their respective locations in the gel. (B) Graphical quantification of intensity of the protein-DNA complex formed.

### 6. Circular Dichroism spectroscopy confirms Myb domain has an alpha helical nature

By analyzing the computational model of the Myb domain, we observed that it was predominantly helical in nature. To confirm the same, circular dichroism (CD) analysis was performed. CD spectroscopy is a well-established technique for elucidating protein secondary structures. It has various applications in structural biology, such as secondary structure determination, protein aggregation, and protein folding. CD spectra in the far-UV region (240–180 nm) corresponded to peptide bond absorption, which could determine the presence of secondary structural elements such as helices, sheets, and turns. CD spectroscopy is frequently employed as a confirmatory method for determining secondary structures because it is quicker and requires minimal resources. The mean residue molar ellipticity, [θ], in degrees cm^2^ dmol^−1^, has been calculated from; [𝜃 ] = [𝜃 ]_obs_× MRW/(10𝑐 𝑙 ) where [θ]_obs_ is the measured molar ellipticity in degrees, MRW is the mean residue molecular weight, *c* is the protein concentration in grams per milliliter, and *l* is the optical path length in centimeters.

CD spectra of the Myb domain were collected in 25 mM Tris buffer (pH 7.5) at 21 ^°^C. The profile shows a clear maximum at 192 nm and two slightly pronounced peaks at 210 nm and 222 nm. After spectral deconvolution (as mentioned in the Materials and Methods section) analysis, the secondary structure estimated the percentages of alpha helices (64.7%), beta sheets (0%), random coils (35.3%), and turns (0%) (average of three runs, Figure 10A and 10B). We also calculated the ratio of ellipticity at 222/208 nm; which yielded a value of 0.94. This value suggests that the protein structure is helical in nature (Andersen et al., 1996). The above results confirmed that the Myb domain is predominantly helical in nature. These results are in agreement with the results of our computational model and the simulation data of the Myb domain.

**Figure 10:**
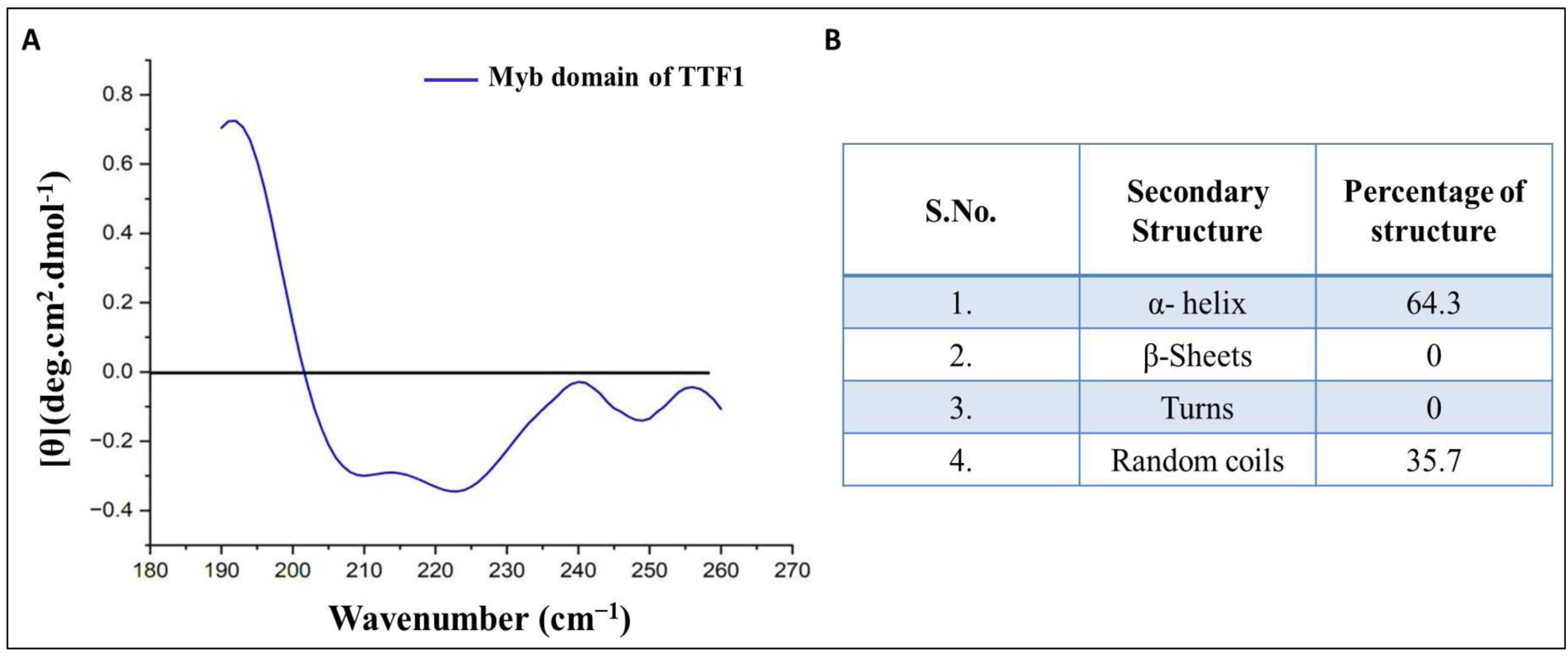
(A) CD spectrum of Myb domain. The x-axis of the spectrum corresponds to the wavenumber, and the y-axis represents molar ellipticity in degrees. (B) Represents the percentage of secondary structures in the Myb domain.

### 7. Raman spectra analysis of Myb domain

Raman spectroscopy is a non-destructive technique that provides comprehensive data on chemical structure, phase and polymorphy, crystallinity, and molecular interactions. The strength and wavelength positions of the scattered Raman light are displayed by the number of peaks in the spectrum. Every peak, including specific bonds like C-C, C=C, N-O, and C-H, is associated with a particular chemical bond vibration. Hence, the secondary structures of proteins and peptides can be effectively derived using Raman spectroscopy. Raman spectral profile is a compilation of signals from various conformations of the protein molecule in solution, offering an "instantaneous snapshot" of the population. Considering above towards characterizing the secondary structure of the purified Myb domain of TTF1 and to comprehend the population heterogeneity, we used Raman spectroscopy. Raman spectrum of the buffer (control) is displayed in black, whereas that of the Myb domain is displayed in red (Figure 11).

**Figure 11:**
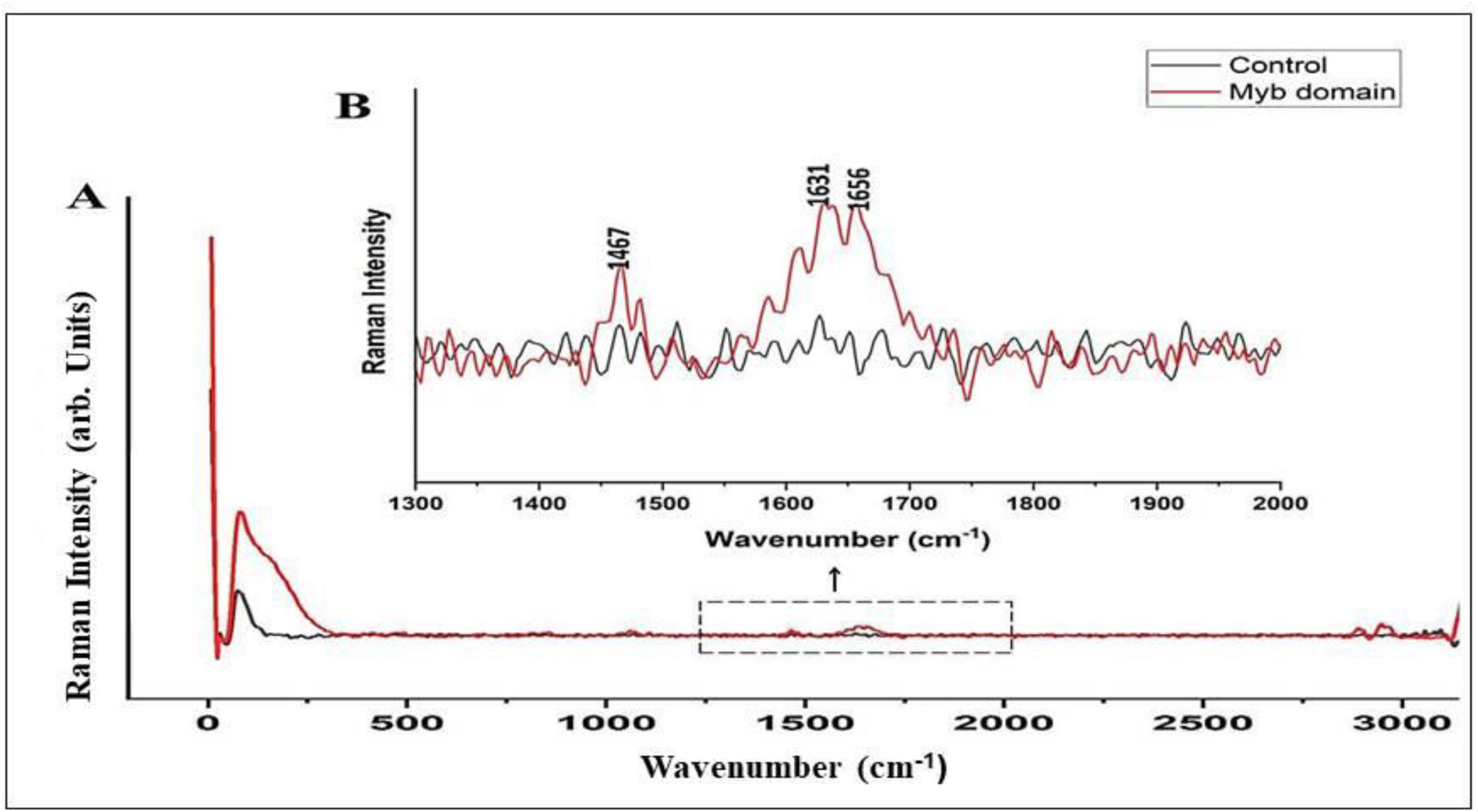
(A) Raman spectrum of the Myb domain of TTF1 protein. (B) Raman spectrum for the 0 –3000 cm^-1^ regions. The Raman peaks were assigned using numerical labels. The Myb domain spectrum is shown in red, whereas that of the buffer control is shown in black. The x axis represents the relative wavenumber (cm^-1^) and y axis represents the Raman intensity (arb. units).

In the spectra, we found various intense peaks which are associated with aromatic amino acids, such as peak at 1000 cm^-1^ is associated with phenylalanine amino acid and that at 1341 cm^-1^ is associated with tryptophan amino acid. The three signals of amide I 1600-1690 cm^-1^ (stretching vibration of C =O), amide II 1480-1580 cm^-1^, and amide III 1230-1300 cm^-1^ (both related to paired C-N stretching and N-H bending vibrations of the peptide group) are of particular importance for the detection of various protein backbone confirmations. Raman spectroscopy of protein secondary structure has adopted the same strategy that is used in analysis. It has concentrated on the relationship between the positions of the amide I and amide III vibrations and the proportion of each secondary structural element in the protein that has been determined through crystallography. The wavenumbers for the amide I and III modes in α-helix and β-sheet structures are typically found to be within the ranges of 1662–1650 and 1272-1264 cm^-1^ (for α-helix) and 1674– 1672 and 1242–1227 cm^-1^ (for β-sheet), respectively. The amide I band, which is mostly formed by C1/4 O stretching with minor contributions, is located within the 1600-1690 cm^-1^ spectral range. The vibrational coupling between the movements of the peptide carbonyl groups arranged in an ordered secondary structure by hydrogen bonds produces a spectrum resulting from the amide I modes. The examination of amide I can provide point-by-point data on the underlying compliance of proteins under physiological circumstances (Figure 11B). We described and examined the amide I spectrum in great detail for the Myb domain because it has the potential to significantly characterize the secondary structure of a protein (Figure 12, Maiti et al., 2004; Signorelli et al., 2017; Tiwari et al., 2022).

**Figure 12:**
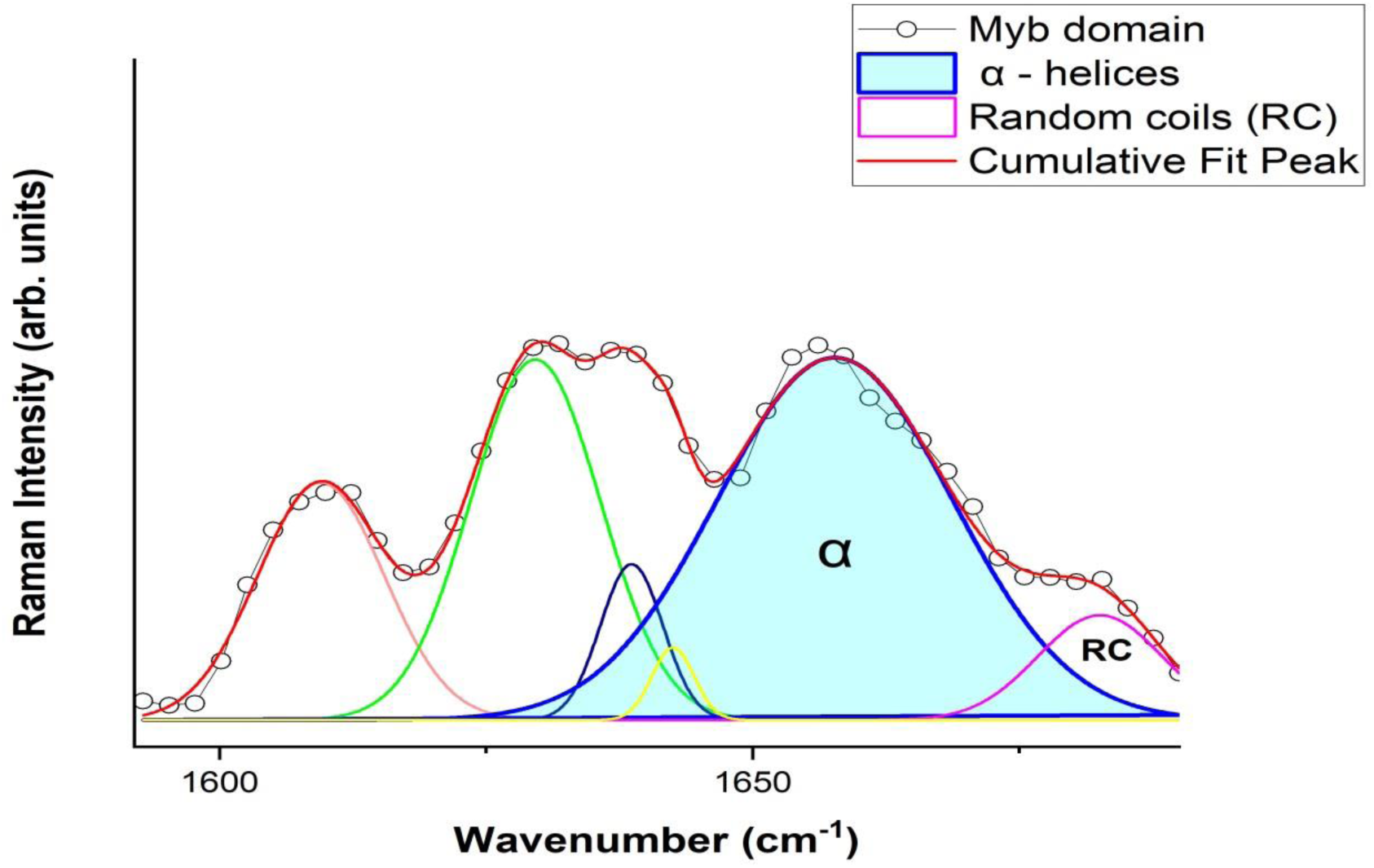
Curve fitting of the amide I (1590 –1690 cm^-1^) region for the Myb domain of TTF1. The x axis represents the relative wavenumber (cm^-1^) and y axis represents the Raman intensity (arb. units).

### 8. Curve fitting for the Amide I region and deconvulation studies

The majority of the C=O stretching was attributed to the amide I region, while the N-H plane deformation component was present in the amide III band. As previously mentioned, our study focused on the amide I peak, as the overlapping side chain vibrations make it challenging to analyze the amide III band. To fully comprehend the secondary structure, we carried out the curve fitting process for the amide I region of the Myb domain. By fitting the measured data into an analytical equation and interpolating between discrete values, we can derive a finite-difference approximation and obtain the maximum or minimum value. The amide I center is approximately 1650 cm^-1^ in α-helical proteins and peptides and functions as a marker band for the ordered α-helix contribution in Raman spectroscopy. More precisely, the band was split into two main parts related to the random coil and alpha helix configurations. Two curves were considered to fit the amide I band: the first at 1650-1656 cm^-1^ was assigned to alpha helices, while the second at 1680 cm^-1^ was linked to random coils. The fitting curves (in amide I) of the Myb domain indicate that the majority fits in the region representing alpha-helices (aqua blue area under the red curve), whereas random coils make up the rest of the curve (Figure 12). These findings align with our computational model and CD spectroscopy data, indicating that the Myb domain is predominantly alpha-helical in nature.

## Discussion

Transcription Termination Factor 1 (TTF1) is a multifunctional nucleolar protein which partakes in several cellular processes, such as Pol I-mediated transcription initiation, termination, r-RNA processing, chromatin remodelling, polar fork arrest, DNA damage, etc. Owing to the importance and multifunctional nature of TTF1, we previously proposed an *ab-initio* model of the full-length mammalian TTF1 protein (Tiwari, et al., 2022). The model revealed that the Myb domain of the protein is a major DNA-binding region. We and others earlier tried to purify both human and mouse full-length TTF1 but were unsuccessful in obtaining homogenous functionally active protein. The full-length protein elutes as aggregates, making it difficult to characterize. As mentioned above, our earlier computational model showed severely disordered regions in the N-terminal domain of the protein (Tiwari, et al., 2022). For the above reasons, we moved ahead towards cloning and characterizing the DNA-binding Myb domain of mouse TTF1. To gain structural insight, we have constructed a 3-D model of the Myb domain via homology and *ab-initio* modelling using different servers such as the Swiss Model, AlphaFold, Robetta, and I-TASSER. The generated model was then analyzed for its stability using Ramachandran analysis and MD simulation, which confirmed that the Robetta provides an excellent model (Figure 3C), as the Ramachandran plot showed very few residues in the prohibited area. The model was comparable to the DNA binding domain structure of the Reb1 protein (crystal structure data, PNAS) and had a very low RMSD value (1.467 nm), proving the good quality of the model. The model reveals four helix turn helix (HTH) motifs in the Myb domain (Figure 3), which is similar to the DNA-binding domain of its homologue Reb-1 (two MybADs and two MybRs). This provided us with the desired confidence in the model, suggesting that we are heading in the right direction towards characterization of this important DNA-interacting domain. Secondary structure prediction shows that the protein is helical in nature (alpha helix ∼65%, Figure 3), which seems to be conserved in Rtf1 of *S. pombe,* Reb1p of *S. cerevisiae,* Nsi1 (Ytt1) of *S. cerevisiae,* and Rib2 of *Xenopus tropicalis* (Jaiswal et al., 2016). To validate the computational model, the Myb domain was cloned into an expression vector and purified to homogeneity (> 95% purity, Figure 8B) through affinity chromatography, followed by gel filtration chromatography. The SDS-PAGE and DLS profiles confirmed the above findings (Figure 8B). Furthermore, the CD spectra and Raman spectroscopy results showed that the Myb domain is a helical protein with approximately 65% alpha helices and 35% random coils (Figure 10A and 10B), which is in agreement with the computational models (Figure 3). Deconvolution of the CD and Raman spectra confirmed that the Myb domain is predominantly helical in nature, which explains its DNA-binding capability. From the existing literature, we can conclude that most of the DNA-binding domains (leucine zippers and HTH Motifs) are helical in nature, which allows DNA to get locked in the major grooves for further activities (Struhl, 1989). Biochemical data (Evers et al., 1995; Grummt et al., 1986) have shown that TTF1 binds to Sal box elements. To check whether the Myb domain alone is sufficient for this activity, we performed an EMSA with the purified protein and Sal box DNA oligos. A shift in the protein and DNA bands confirmed the successful formation of this complex. The above results prove that the Myb domain alone is sufficient for DNA binding. As has been observed for other transcription factors, various domains are involved in different regulatory functions.

Hence, our work is the first to characterize this essential domain structurally, opening up possibilities for further investigation. Earlier, obtaining the homogeneously purified functional TTF1 protein was a rate-limiting step towards solving the atomic structure of this essential protein. Since we have purified the functional Myb domain of TTF1, we are moving ahead towards solving the atomic structure of this protein either by crystallography or NMR. Solving the structure will provide insights into the mechanism by which this domain engages DNA for various activities. Also, our lab is ambitiously exploring the possibilities to purify the full-length protein in order to understand the regulatory mechanism of this essential transcription factor, which is also engaged in regulating DNA replication and DNA damage repair.

## Conclusion

Conclusively, this is the first ever study to report a complete *in silico* model of the Myb domain of the mouse TTF1 protein validated by experimental procedures after purifying the protein to homogeneity. It is concluded that the Myb domain contains ∼65% alpha helices and on its own able to bind DNA.

## Acknowledgement

The authors are thankful to the Director Prof. Sanjay Kumar, Institute of Science, Banaras Hindu University for providing space and facilities to conduct the research. We are also thankful to the Central Discovery Centre (CDC) and Sophisticated Analytical and Technical Help Institutes (SATHI), Banaras Hindu University (BHU), for the accessibility of the CD spectroscopy and Raman spectroscopy instruments. We are also grateful to the support and the resources provided by ‘PARAM Shivay Facility’ under the National Supercomputing Mission, Government of India at the Indian Institute of Technology (BHU), Varanasi. The research was funded by the Department of Biotechnology (DBT), Govt. of India, RLS grant (BT/RLF/Re-entry/43/2016) and IOE seed grant to Samarendra K Singh. The Council of Scientific and Industrial Research (CSIR) supported this research by providing scholarship to Gajender Singh, Department of Biotechnology (DBT), and Govt. of India by providing scholarship to Saloni Khatri and Prashant Prakash (M.Sc. Stipend). We further show our thanks to ISLS for providing space and instrumentation support.

## Author contribution

SKS was involved in the conceptualization and designing; supervision; writing - critical review & editing of the manuscript. SMS was involved in supervision and original manuscript writing. GS was involved in investigation, methodology data collection, visualization & analysis and writing, review & editing of the manuscript. SK and PP helped GS in data analysis and manuscript writing. AJB and RK helped in MD simulation design, analysis, writing and review of the manuscript.

## Supplementary information

#### Box 1

Amino acid sequence of Myb domain of TTF1

Q62187 Uniprot ID of mouse TTF1

Myb Domain: 550 to 732 amino acids

Sequence:

TLITNLKRKHAFRLHIGKGIARPWKLVYYRAKKIFDVNNYKGRYNEEDTKKLKAYHSLHGNDWKKIGAMVARSSLSVALKFSQIGGTRNQGAWSKAETQRLIKAVEDVILKKMSPQELRELDSKLQEDPEGRLSIVREKLYKGISVEARVETRNWMQCKSKWTEILTKRMTHGGFVYRGV

**Figure S1:**
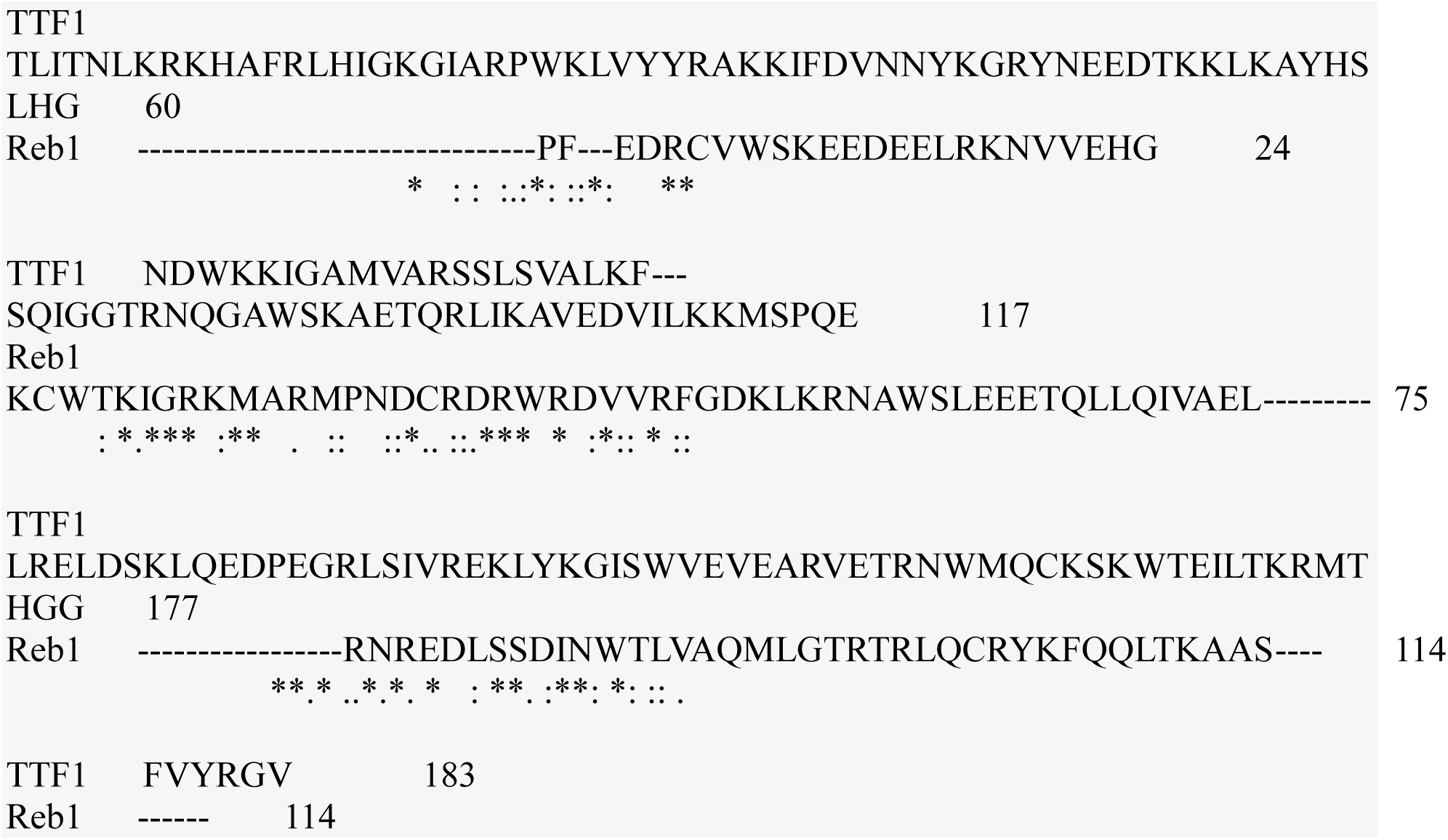
Sequence alignment of the Myb domain of mouse TTF1 and Reb1 protein

**Figure S2:**
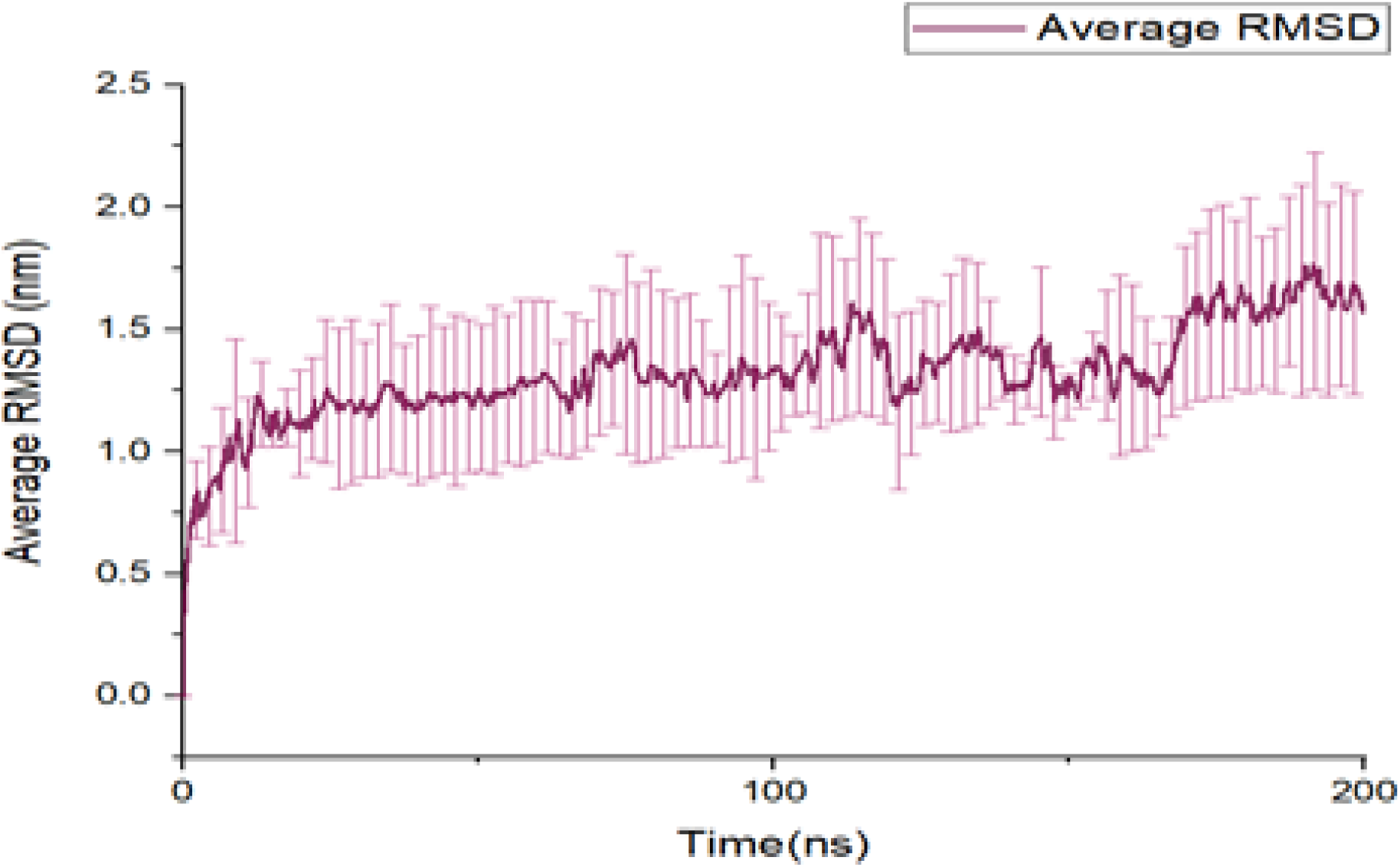
Average of RMSD value for triplicate run.

**Figure S3:**
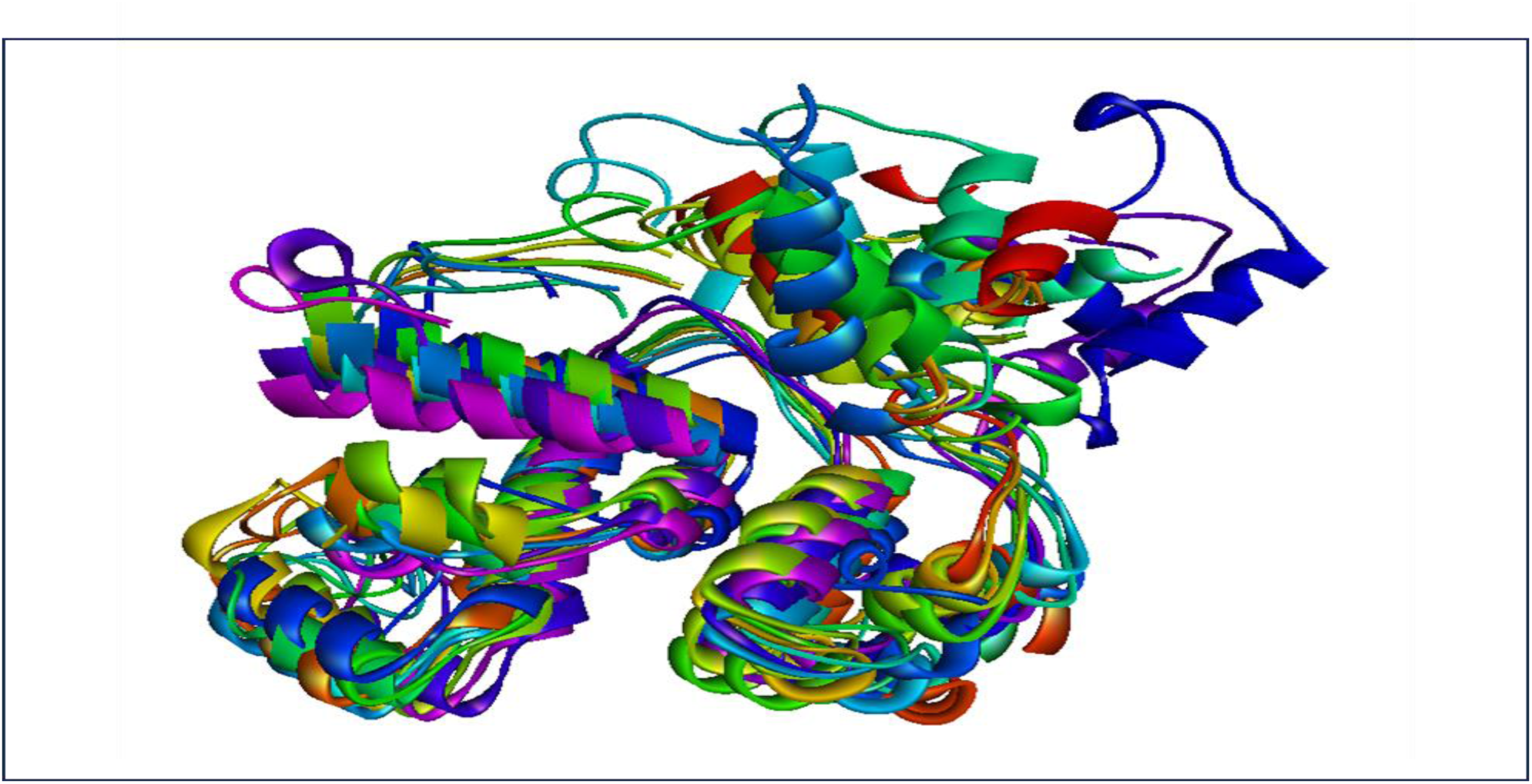
The overlay of protein structure after every 20 ns. Frame for protein after every 20 ns were extracted and align using PyMOL

**Figure S4:**
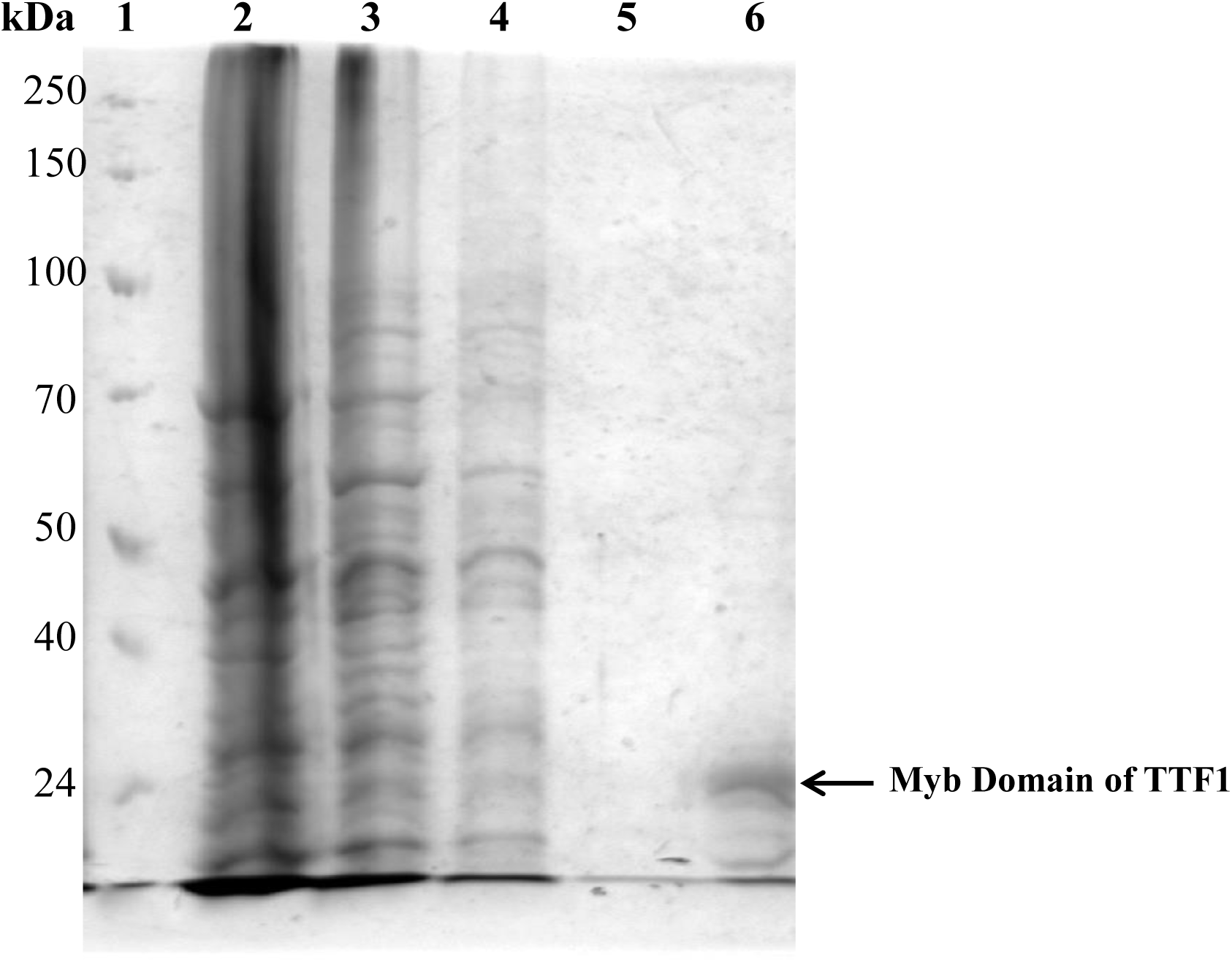
SDS-PAGE profile of the affinity purified protein. Lane 1 protein marker, Lane 2 Cell lysate, Lane 3 Flow through, Lane 4 &5 Wash 1 and Wash 2, Lane 6 Elution 1.

